# Curcumin-mediated photodynamic inactivation of *Escherichia coli, Pseudomonas fluorescens*, and *Candida auris*

**DOI:** 10.1101/2025.08.18.670777

**Authors:** Jaime A. Garcia-Diosa, Bhanu K. Pothineni, Adrian Keller

**Affiliations:** Paderborn University, Technical and Macromolecular Chemistry, Warburger Str. 100, 33098 Paderborn, Germany

**Keywords:** curcumin, photosensitizer, antimicrobial photodynamic therapy, bacteria, fungi

## Abstract

Photodynamic therapy (PDT) is an emerging antimicrobial strategy that uses light to activate photosensitizers, generating reactive oxygen species (ROS) that induce microbial cell death. This study investigates the antimicrobial efficacy of curcumin-mediated PDT against the microbial pathogens *Escherichia coli, Pseudomonas fluorescens*, and *Candida auris*. The minimum inhibitory concentration (MIC) for each microorganism is determined at different light fluence rates within a clinically relevant range. Our results demonstrate that curcumin-mediated PDT effectively inactivates *E. coli* and *C. auris*, while exhibiting only partial inhibition against *P. fluorescens* at curcumin concentrations up to 60 µM. Notably, within the tested fluence rate range, the total light dose appears to be a more critical determinant of antimicrobial efficacy than the specific fluence rate itself. The observed variations in microbial susceptibility highlight the importance of species-specific factors, such as cell wall/membrane structure. These findings provide guidelines for the application of curcumin-mediated PDT in combatting antimicrobial resistance.

## 1. Introduction

The escalating challenge of antimicrobial resistance (AMR) calls for novel therapeutic strategies to combat infections caused by bacteria, fungi, viruses, and parasites (Laxminarayan et al. 2016). Traditional antibiotic treatments are becoming increasingly ineffective, posing a significant threat to global health. In this context, photodynamic therapy (PDT) has emerged as a promising therapeutic modality that utilizes light to activate photosensitizing agents, generating reactive oxygen species (ROS) that damage essential cellular components and ultimately cause the death of the microorganism (Dougherty et al. 1998; Ghorbani et al. 2018). PDT has demonstrated efficacy against a broad spectrum of microorganisms, including multidrug-resistant bacterial and fungal strains, positioning it as a valuable tool in combating AMR (Youf et al. 2021). Unlike conventional antibiotics, the multi-target mechanism of action of PDT reduces the likelihood of resistance development (Wainwright 1998; Youf et al. 2021). Resistance to antimicrobial PDT (aPDT) is rarely reported, suggesting that while the possibility of microbial adaptation and escape from this treatment may exists, it is currently limited (Cieplik et al. 2018). This is a crucial advantage in the face of rapidly evolving resistance mechanisms in pathogenic bacteria and fungi.

Despite its advantages, aPDT faces a number of challenges that impede its widespread clinical adoption. Common photosensitizers (PSs) often suffer from issues such as limited tissue penetration, lack of selectivity, aggregation and quenching, and dark toxicity, resulting in intense research efforts dedicated to the development and discovery of new PSs (Klausen et al. 2020; Polat and Kang 2021; Huis In ‘t Veld et al. 2023). Curcumin, a naturally occurring polyphenol derived from turmeric (*Curcuma longa*), has emerged as a promising alternative due to several key advantages. These include high biocompatibility, low systemic toxicity, and synergistic therapeutic effects, such as anti-proliferative, anti-inflammatory, and anti-angiogenic activities (Dias et al. 2020). Consequently, it is investigated in various implementations of PDT, most importantly in cancer therapy (Xie et al. 2022) but also in the treatment of oral infections (Santezi et al. 2018). Furthermore, since curcumin is a common food ingredient and food additive, it is frequently used in the photodynamic inactivation (PDI) of foodborne pathogens in various food products (Yu et al. 2022; Lan et al. 2023).

The PDT effect depends on the interaction of three essential components: molecular oxygen, the PS, and light. The PDT dose can be controlled by adjusting the latter two components. It is generally assumed that the biological effect of light in PDT is determined by the total light dose, regardless of the exposure time (Gao et al. 2020). Consequently, PDT studies often present data in terms of total light dose to compare the PDI of microorganisms (Dias et al. 2020; Jao et al. 2023; Law et al. 2023). However, the fluence rate, *i.e.*, the power density of light, may influence PDT efficacy by affecting metabolic oxygen consumption and, consequently, the efficiency of ROS production (Nichols and Foster 1994; Henderson et al. 2006; Ghorbani et al. 2018). Indeed, some studies on the photodynamic treatment of cancer cells and tumors have shown that lower fluence rates can enhance the cytotoxic effect of the treatment by optimizing the depletion of the cell’s antioxidant defenses (Angell-Petersen et al. 2006; Yamamoto et al. 2006; Wang et al. 2007; Woodhams et al. 2007; Grossman et al. 2016; Dos Santos et al. 2020). However, there are only few reports on this phenomenon in aPDT. Qin *et al.* conducted a study on the effect of different parameters of toluidine blue-mediated PDI effectiveness, including fluence rate, on the PDI of periodontal pathogens. They observed that a high fluence rate (159 mW cm^-2^) increases the cytotoxicity of the treatment (Qin et al. 2008). A similar conclusion was reported by Prates *et al.*, who studied the effects of fluence rate and light dose on the treatment of three different strains of *Candida sp.* using methylene blue as the PS. In these experiments, a higher fluence rate (300 mW cm^-2^) lead to a higher cytotoxicity of the treatment, although the production of ROS is inversely proportional to the fluence rate (Prates et al. 2009). Due to the limited number of studies, the role of the fluence rate in aPDT is not fully understood. Additionally, these studies used fluence rates that are much higher than those recommended to avoid possible damage to host tissue in clinical applications (Sitnik and Henderson 1998; Jori et al. 2006).

In this study, we evaluate the efficacy of curcumin-mediated PDI of three relevant pathogens by determining their minimum inhibitory concentrations (MICs) at different fluence rates within a clinically relevant range. It has been observed before that different bacterial species show markedly different susceptibilities to curcumin-mediated PDI (Penha et al. 2017), which have to be considered in the development of effective treatment protocols. For the current investigation, we have selected *Escherichia coli, Pseudomonas fluorescens*, and *Candida auris* as the target pathogens. *E. coli* is a Gram-negative bacterium that causes a wide range of infections (Nataro and Kaper 1998; Rangel et al. 2005) and thus is a frequent target of aPDT. *P. fluorescens* is a common food contaminant capable of growing at a wide range of temperatures, making it a significant issue in both refrigerated and unrefrigerated food items (Kumar et al. 2019). Curcumin-mediated PDI of *P. fluorescens*, therefore, has been investigated almost exclusively in food items (Saraiva et al. 2021; Saraiva et al. 2024; Zheng et al. 2025). However, *P. fluorescens* can also cause severe infections in immunocompromised individuals, most importantly bacteremia due to contaminated medical equipment (Gershman et al. 2008), but also skin and soft tissue infections, as well as respiratory tract infections (Liu et al. 2021), which warrants investigations also in the context of aPDT settings. *C. auris* is a newly emerged fungal pathogen that has become a global threat due to its rapid spread and multidrug resistance (Du et al. 2020). PDI of *C. auris* has already been demonstrated using several PSs (Bapat and Nobile 2021; Grizante Barião et al. 2022; Silva et al. 2023), but not with curcumin. By investigating the combined effects of curcumin concentration and fluence rate on these target pathogens, this study aims to contribute to a better understanding of the factors influencing the efficacy of curcumin-mediated aPDT and its potential as a novel antimicrobial strategy to combat the growing challenge of antimicrobial resistance.

## 2. Results

Figure 1 evaluates *E. coli* growth after 420 nm irradiation with a fluence rate of 17.8 mW cm^-2^ and 76.3 mW cm^-2^, respectively. The control samples incubated in the dark exhibit a relatively stable OD_600_ value of approximately 0.55, indicative of normal bacterial growth and no dark cytotoxicity (see Figure 1A). In contrast, the samples irradiated at 420 nm with both fluence rates (17.8 mW cm^-2^ and 76.3 mW cm^-2^) demonstrate a sharp decline from OD_600_ values from about 0.55 to about 0.15 at curcumin concentrations of 7.5 µM and above (see Figure 1A). This low OD_600_ value corresponds to that of the negative control (see Figure S1), indicating complete growth inhibition. Notably, no significant difference in bacterial growth inhibition is observed between the two fluence rates. This suggests that the maximum PDI effect is reached once the curcumin concentration is 7.5 µM or above, regardless of the fluence rate within the therapeutic window. The images of 24 h *E. coli* cultures on agar plates after PDI provide a visual confirmation of the quantitative OD_600_ data (see Figure 1B and Figure S2). The control plates (black outline) show confluent bacteria at the maximum curcumin concentration investigated, consistent with the OD_600_ values. After PDI, plates corresponding to curcumin concentrations of 1.88 µM and 7.5 µM (blue and red outlines) show a clear reduction in colony forming units compared to the control, consistent with the OD_600_ measurements.

**Figure 1.**
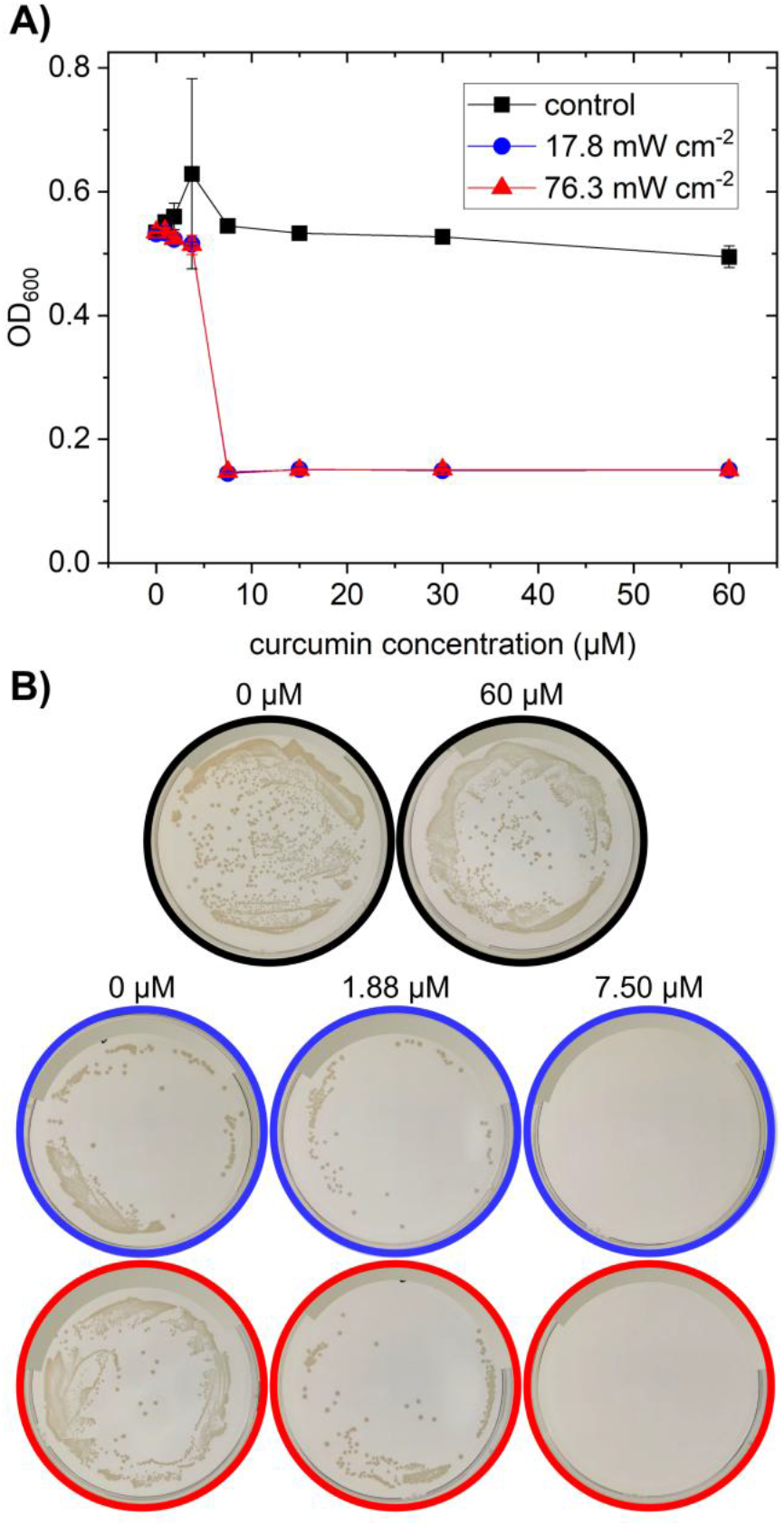
*E. coli* growth after curcumin-mediated PDI using 420 nm light at a dose of 20 J cm^-2^ with fluence rate of 17.8 mW cm^-2^ and 76.3 mW cm^-2^, respectively, at different curcumin concentrations. A) Optical density (OD_600_) after 24 h of incubation at 37 °C. Error bars indicate the standard deviation (n = 3). Controls were incubated in the dark. See Figure S1 for initial OD_600_ values (0 h) and negative controls. B) *E. coli* cultures (24 h) on agar after PDI at different curcumin concentrations and fluence rates. Control (incubated in the dark, black outline), low fluence rate (blue outline), and high fluence rate (red outline). See Figure S2 for all plates.

Figure 2 presents the results of curcumin-mediated PDI on *P. fluorescens* with both fluence rates at different curcumin concentrations. Again, the control samples exhibit a stable OD_600_ value, indicating consistent bacterial growth and no dark cytotoxicity. In contrast, samples treated with PDI demonstrate a concentration-dependent decrease of OD_600_ values, indicative of a reduction of bacterial viability (see Figure 2A). At the lower fluence rate (17.8 mW cm^-2^), a remarkable linear decrease of OD_600_ is observed at curcumin concentration higher than 15 µM. Similarly, at the highest fluence rate (76.3 mW cm^-2^), OD_600_ values decrease with increasing curcumin concentration. However, this decrease starts already at the lowest curcumin concentration investigated and appears to saturate for the highest concentrations. Bacterial growth is detected even at 60 µM curcumin. This is consistent with the *P. fluorescens* cultures on agar plates shown in Figure 2B and Figure S4. Samples after PDI at 15 µM and 60 µM curcumin at both fluence rates (blue and red outlines), show a clear reduction in colony forming units compared to the control, but no complete bacterial inhibition is observed.

**Figure 2.**
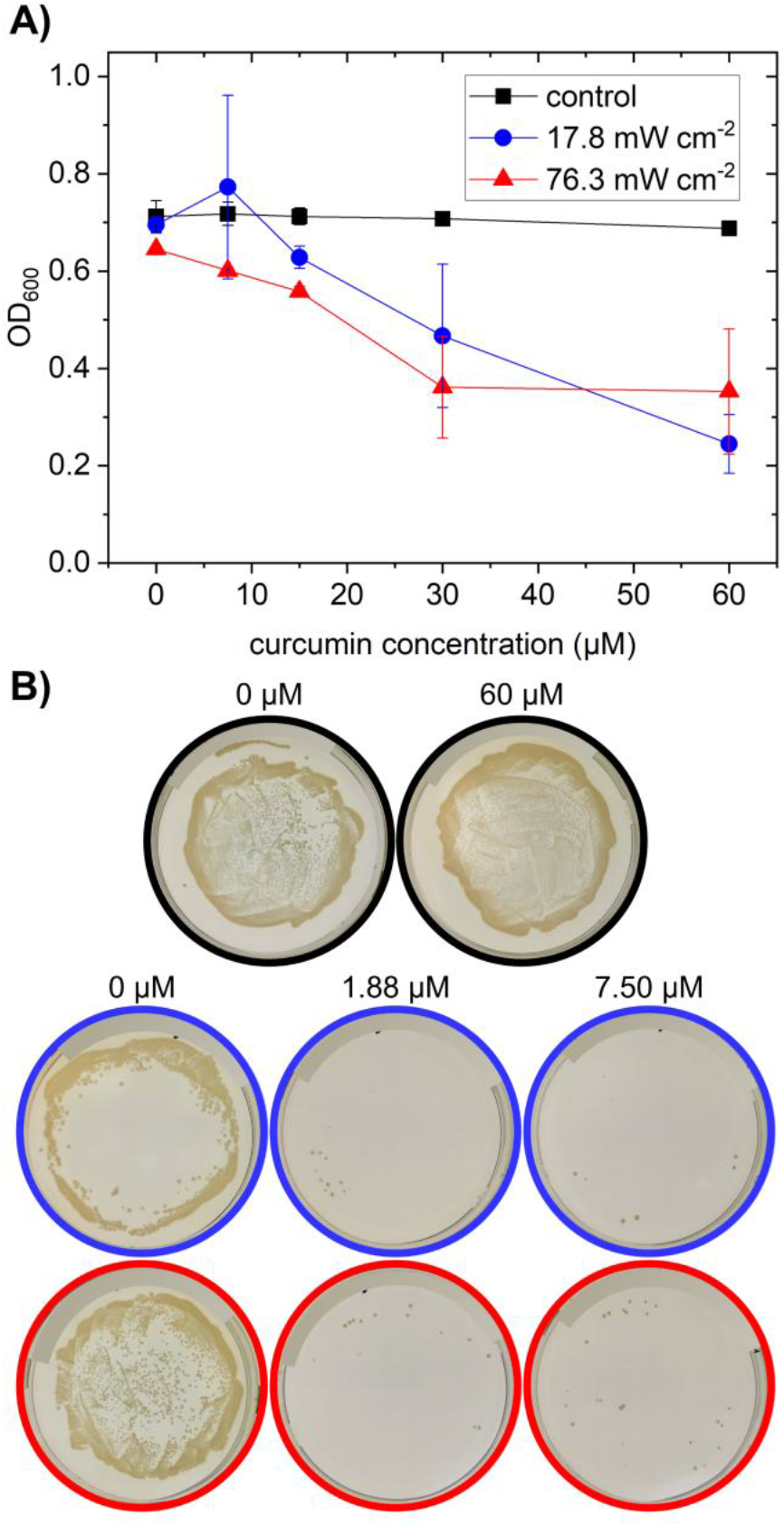
*P. fluorescens* growth after curcumin-mediated PDI using 420 nm light at a dose of 20 J cm^-2^ with fluence rate of 17.8 mW cm^-2^ and 76.3 mW cm^-2^, respectively, at different curcumin concentrations. A) Optical density (OD_600_) after 24 h of incubation at 30 °C. Error bars indicate the standard deviation (n = 3). Controls were incubated in the dark. See Figure S3 for initial OD_600_ values (0 h) and negative controls. B) *P. fluorescens* cultures (48 h) on agar after PDI at different curcumin concentrations and fluence rates. Control (incubated in the dark, black outline), low fluence rate (blue outline), and high fluence rate (red outline). See Figure S4 for all plates.

The results of curcumin-mediated PDI on *C. auris* at low and high fluence rates are shown in Figure 3. The control samples exhibit a relatively stable OD_600_ value around 0.8 – 0.9 across varying curcumin concentrations, showing consistent growth of the yeast and no dark cytotoxicity (see Figure 3A). On the other hand, irradiated samples show a markedly concentration-dependent decrease in OD_600_ values. At both fluence rates (17.8 mW cm^-2^ and 76.3 mW cm^-2^), a strong reduction of the fungal growth is observed even at the lowest curcumin concentration used (1.87 µM). When the curcumin concentration reaches 15 µM and more, the OD_600_ values approaches 0.15, corresponding to the OD_600_ of the negative control (see Figure S5), indicating a near-complete inhibition of fungal growth (see Figure 3A). Notably, there is again no significant difference between the two fluence rates at any of the tested curcumin concentrations. This suggests that within the tested fluence range, the curcumin concentration is the primary parameter of the photodynamic effect. *C. auris* cultures on agar plates after PDI visually confirm the trends observed in the OD_600_ measurements (see Figure 3B and Figure S3). The control group shows dense *C. auris* growth, consistent with the high OD_600_ values. In the case of the samples irradiated at both fluence rates (blue and red outlines), a strong reduction in the number of colonies is observed at 3.75 µM, whereas at 15 µM, visible colonies are almost complete absent (see Figure 3B and Figure S6), corroborating the trend of the OD_600_ values.

**Figure 3.**
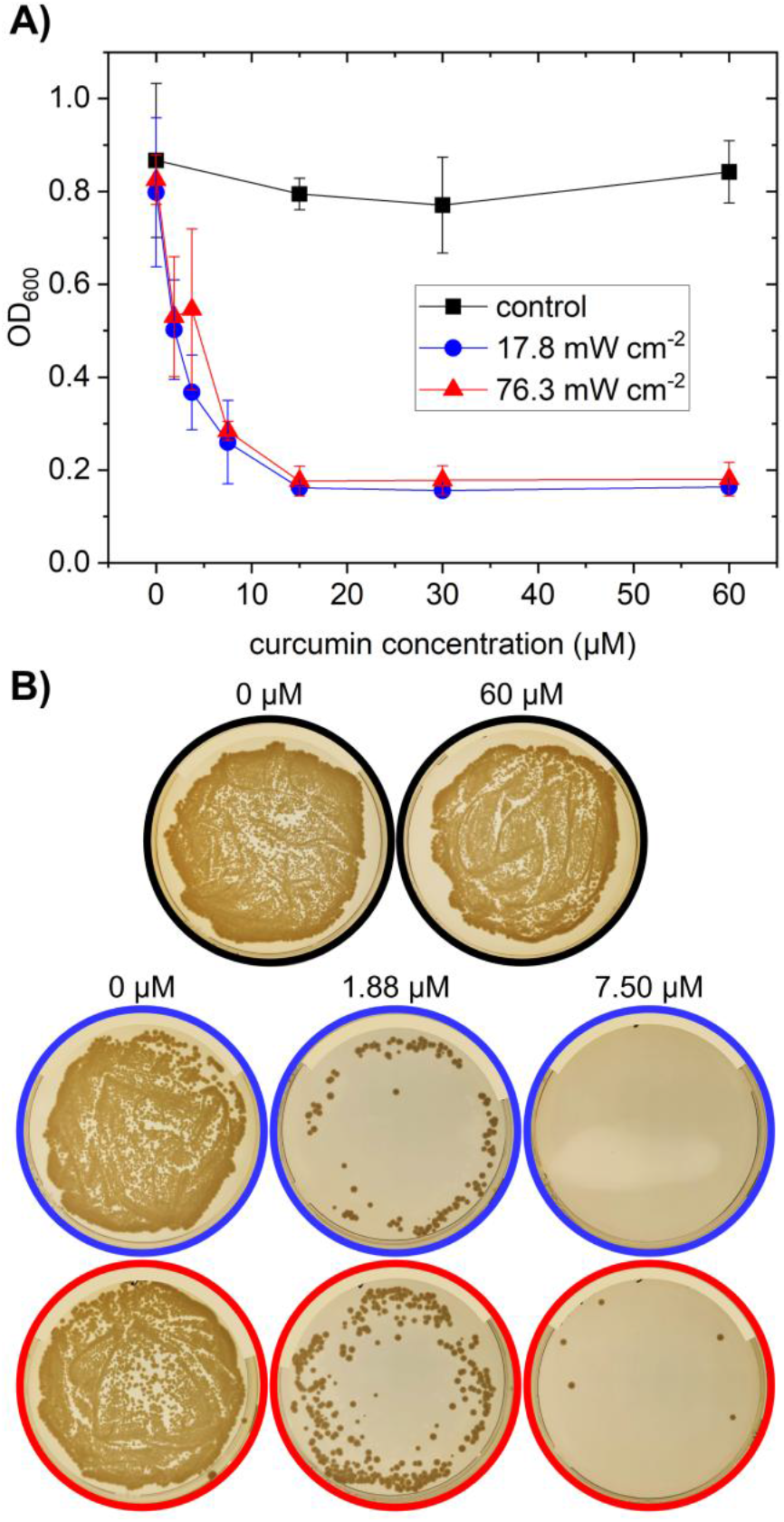
*C. auris* growth after curcumin-mediated PDI using 420 nm light at a dose of 20 J cm^-2^ using with fluence rate of 17.8 mW cm^-2^ and 76.3 mW cm^-2^, respectively, at different curcumin concentrations. A) Optical density (OD_600_) after 24 h of incubation at 37 °C. Error bars indicate the standard deviation (n = 3). Controls were incubated in the dark. See Figure S5 for initial OD_600_ values (0 h) and negative controls. B) *C. auris* cultures (24 h) on agar after PDI at different curcumin concentrations and fluence rates. Control (incubated in the dark, black outline), low fluence rate (blue outline), and high fluence rate (red outline). See Figure S6 for all plates.

## 3. Discussion

Table 1 summarizes the MIC of curcumin during aPDT for *E. coli, P. fluorescens*, and *C. auris* as determined from the OD_600_ results. The data reveal a remarkable difference in the susceptibility of the three microorganisms to curcumin-mediated aPDT. *E. coli* exhibits the highest sensitivity of the treatment, with a MIC of 7.5 µM, implying that a relatively low concentration of curcumin is sufficient to inhibit *E. coli* growth. *C. auris* demonstrates intermediate susceptibility with a MIC of 15 µM, requiring a higher curcumin concentration to effectively inhibit the growth of the yeast. On the other hand, *P. fluorescens* shows the lowest susceptibility to curcumin-mediated aPDT with a MIC greater than 60 µM, indicating that *P. fluorescens* is significantly more resistant to the photodynamic effects of the PS curcumin under the studied conditions.

**Table 1.** Minimum inhibitory concentrations (MICs) of curcumin during aPDT using a fluence rate range of 17.8 – 76.3 mW cm^-2^, and a total light dose of 20 J cm^-2^.

| Microorganism | MIC (µM) |
| --- | --- |
| *E. coli* | 7.5 |
| *P. fluorescens* | > 60.0 |
| *C. auris* | 15.0 |

Our results demonstrate that curcumin-mediated aPDT is effective in inactivating *E. coli* and *C. auris*, and partially inactivating *P. fluorescens* at curcumin concentrations up to 60 µM. These results are consistent with previous studies, which have reported varying degrees of susceptibility to curcumin-mediated aPDT among different microorganisms (Dovigo, Pavarina and Ribeiro et al. 2011; Andrade et al. 2013; Penha et al. 2017; Saraiva et al. 2021; Saraiva et al. 2024). For instance, *P. fluorescens* has been shown to be more resistant to curcumin-mediated aPDT compared to *E. coli* and *C. albicans* (Saraiva et al. 2021; Saraiva et al. 2024). This difference in susceptibility may be attributed to several factors, including variations in cell wall/membrane structure and composition, and variations in the uptake and intracellular accumulation of curcumin among the different microorganisms. In general, curcumin can interact with microorganisms through several mechanisms, including charge interactions, which are pH-dependent due to keto-enol tautomerism (Priyadarsini 2009), binding to various enzymes such as sortase A (Hu et al. 2013; Praditya et al. 2019) and FtsZ (Kaur et al. 2010), and membrane adsorption and subsequent uptake into the cell. Species-specific differences may play a role in all these mechanisms but are particularly important for cellular uptake (Annunzio et al. 2018).

This study highlights the importance of species-specific variations in susceptibility to curcumin-mediated aPDT. Despite sharing a common Gram-negative cell wall structure with an outer membrane containing lipopolysaccharides (LPSs), *E. coli* and *P. fluorescens* display markedly different responses to aPDT with curcumin (see Table 1). While the precise mechanisms governing the interaction of curcumin with these species remain to be fully elucidated, the high efficacy of curcumin-mediated aPDT against *E. coli* suggests that curcumin can effectively penetrate the outer membrane, likely via charge-mediated interactions, leading to subsequent membrane disruption by the generated ROS (Oliveira et al. 2018; Gao et al. 2019). In contrast, the significantly elevated MIC for *P. fluorescens* suggests a potential limitation in curcumin accumulation within the cell, probably due to reduced passive uptake.

In the case of *C. auris*, a cell wall composed of chitin, glucans, and mannans is present, which differs significantly from bacterial cell walls (Free 2013). However, the intermediate susceptibility to curcumin-mediated PDI compared to *E. coli* and *P. fluorescens* (see Table 1), suggests that curcumin can effectively penetrate the *C. auris* cell wall and accumulate within the cell at a higher rate than for *P. fluorescens*. This is in line with previous observations of effective curcumin-mediated PDI of various *Candida* species (Dovigo, Pavarina and Carmello et al. 2011; Dovigo, Pavarina and Ribeiro et al. 2011; Andrade et al. 2013). The observed variations in susceptibility among the different microorganisms underscore the importance of considering species-specific factors, such as cell wall/membrane structure, and curcumin uptake mechanisms, when evaluating the effectiveness of aPDT.

Regarding the fluence rate effect on the PDT efficiency, several studies in cancer treatment suggest that lower fluence rates at the same total light dose with a longer treatment time can be more efficient in PDT compared to higher fluence rates (Henderson et al. 2006; Dos Santos et al. 2020; Gao et al. 2020). The reasons behind this effect of the fluence rate are still under investigation, but some studies suggest that oxygen availability plays an important role. Lower fluence rates would allow for more efficient oxygen diffusion into the target tissue, whereas high fluence rates lead to the depletion of available oxygen, thereby limiting ROS generation and ultimately reducing PDT efficacy (Dougherty et al. 1998; Alvarez and Sevilla 2024). Conversely, some reports suggest an inverse relationship between fluence rate and aPDT efficiency (Qin et al. 2008; Prates et al. 2009). While higher fluence rates may lead to increased PS bleaching, they have also been associated with greater microbial inhibition (Prates et al. 2009). This apparent contradiction may be explained by a potential initial surge in ROS formation at higher fluence rates, followed by photobleaching. However, increasing the fluence rate infinitely to enhance aPDT efficacy is not feasible. Photon absorption by the PS is limited, restricting ROS production, and the photobleaching effect becomes increasingly significant (Qin et al. 2008). Furthermore, to minimize host tissue damage, the therapeutic window for aPDT is generally considered to be below approximately 50 mW cm^-2^ (Jori et al. 2006).

In this study, the tested fluence rates (17.8 – 76.3 mW cm^-2^) did not significantly influence the antimicrobial activity within the tested concentration range. This finding suggests that the moderate changes of the fluence rate possible within the therapeutic window are insufficient to induce significant variations in ROS generation. Under these conditions, the total light dose thus appears to be the primary determinant of aPDT efficacy. Nevertheless, future studies should investigate a wider range of fluence rates and total light doses to further elucidate the impact of these parameters on aPDT efficacy. Furthermore, investigating the underlying mechanisms of curcumin-mediated PDI in these and other microorganisms, including the role of specific cellular targets and the generation of ROS, will provide valuable insights for optimizing this approach and developing novel antimicrobial strategies.

## 4. Experimental

### 4.1. Bacteria culture

*E. coli* W1485 (DSM 5695) and *P. fluorescens* 1175 (DMS 1976) were purchased from Leibniz Institute DSMZ. For bacteria culture preparation, two or three bacterial colonies were suspended in 30 mL LB medium (MP Biomedicals) and incubated overnight at 37 °C for *E. coli* or 24 h at 30 °C for *P. fluorescens*. In the case of *E. coli*, 250 μL overnight suspension was used to inoculate fresh LB medium and incubated for 4-6 h at 37 °C. For *P. fluorescens*, the culture incubated for 24 h was used directly. Lastly, the bacteria suspensions were centrifuged, washed in 5 % dimethyl sulfoxide (DMSO, Carl Roth), and resuspended in 5 % DMSO, adjusting the optical density at 600 nm (OD_600_) to 0.2. DMSO was used as a solvent to ensure the solubility of curcumin (Saraiva et al. 2021). At a concentration of 5 %, DMSO has been reported to exert minimal cytotoxic effects on bacteria (Ansel et al. 1969; Tunçer Çağlayan and Gurbanov 2024). DMSO solutions were prepared in water instead of buffer because the presence of ions may affect the efficiency of PDI (Núñez et al. 2014). While the resulting osmotic shock might affect bacterial growth to some extent, bacteria are able to resume their original growth rate within tens of minutes (Rojas et al. 2014). Furthermore, samples and controls were treated in the same way, so that possible residual effects will not influence the interpretation of the results.

### 4.2. Fungi culture

*C. auris* (DSM 105990) was purchased from Leibniz Institute DSMZ. The fungi culture was prepared using two or three colonies, which were resuspended in 30 mL yeast extract peptone dextrose medium (YPD medium, SERVA) and incubated at 180 rpm and 37 °C overnight. Finally, the culture was centrifuged, washed in 5 % DMSO, and resuspended in 5 % DMSO, adjusting the optical density (OD_600_) to 0.2.

### 4.3. Antimicrobial photodynamic inactivation

PDI was performed using a Lumidox II 96-position LED array 420 nm with cooling base (Analytical Sales and Services) with Curcumin (Sigma Aldrich) as the PS. The experiments were performed in triplicate using the microdilution method in sterile 96-wells microplates. 50 μL aliquots of microbial suspension in 5 % DMSO (OD_600_ = 0.2) were added to 50 μL of curcumin in 5 % DMSO. The microplates were incubated for 30 min at 25 °C before PDI. Control samples without curcumin and light irradiation were also prepared to analyze the curcumin toxicity and light-induced inhibition.

The fluence rate effect in the PDI was evaluated at a constant light dose of 20 J cm^-2^ using the lowest and highest fluence rate levels available in the Lumidox II equipment. Since the 96-well microplate was held at 17 mm from the light source, the samples were irradiated for 1123 s at the lowest fluence rate (17.8 mW cm^-2^), and for 262 s at highest fluence rate (76.3 mW cm^-2^), obtaining a light dose of 20 J cm^-2^ at both fluence rates. At these short exposure times, the potential impact of prolonged DMSO exposure on bacterial growth is minimized. For subsequent minimum inhibitory concentration (MIC) assays, the DMSO concentration was reduced to 1 %, a level generally considered to have negligible or no effect on bacterial viability (Ansel et al. 1969; Liu et al. 2023). Furthermore, all PDI experiments were conducted in the absence of bacterial growth media and ions to prevent light absorption by medium components, and to minimize interactions with ROS, which could ultimately reduce the effectiveness of PDI.

### 4.4. Optical density measurements

PDI at the different fluence rates was studied by measuring the OD_600_ of the treated samples after 24 h of incubation in their respective media. After treatment, a 20 μL aliquot of each sample was added to 80 μL (1 % DMSO final concentration) of LB (*E. coli and P. fluorescens*) or YPD medium (*C. auris*), in sterile 96-well microplates. Then, the microplates were incubated at 37 °C (*E. coli* and *C. auris*) or at 30 °C (*P. fluorescens*). The OD_600_ values were measured at the beginning and after 24 h of incubation using a microplate reader TrisStar^2^ S LB 942 (Berthold technologies).

### 4.5. Agar plate culture

For visual inspection of PDI, 80 μL aliquots of treated samples were spread on agar plates and incubated. *E. coli* was incubated on LB agar plates at 37 °C for 24 h. *P. fluorescens* was incubated on LB agar plates at 30 °C for 48 h. *C. auris* was incubated on YPD agar plates at 37 °C for 24 h.

### 4.6. Statistical analysis

The OD_600_ was measured in triplicate for each experimental condition (n = 3). Data are presented as mean ± standard deviation (SD) of the three values.

## 5. Conclusions

This study investigated the antimicrobial efficacy of curcumin-mediated PDI against *E. coli, P. fluorescens*, and *C. auris* using a range of curcumin concentrations and two distinct fluence rates. Our findings demonstrate that curcumin-mediated aPDT is able to effectively inactivate *E. coli* and *C. auris*, while leading only to partial inhibition of *P. fluorescens* at curcumin concentrations up to 60 µM. Notably, within the tested fluence rate range (17.8 – 76.3 mW cm^-2^), no significant difference in antimicrobial activity was observed, suggesting that within the therapeutic window, the total light dose is a more critical determinant of aPDT efficacy than the specific fluence rate itself.

The observed variations in microbial susceptibility to curcumin-mediated PDI highlight the importance of species-specific factors, such as cell wall/membrane structure, and curcumin uptake and accumulation. While the mechanisms responsible for the observed differences remain to be fully elucidated, our results suggest that curcumin effectively penetrates the bacterial cell wall of *E. coli* and the fungal cell wall of *C. auris*, leading to significant growth inhibition. In contrast, the limited efficacy against *P. fluorescens* may be attributed to reduced curcumin uptake or increased efflux mechanisms.

These findings provide valuable insights into the potential of curcumin-mediated aPDT as an antimicrobial strategy. However, further research is warranted in order to optimize treatment parameters, including a comprehensive evaluation of the effects of different total light doses and a wider range of fluence rates. Additionally, investigating the underlying mechanisms of curcumin-mediated PDI in these microorganisms, including the specific cellular targets and the generation of ROS, will be crucial for developing more effective and targeted antimicrobial therapies.

## Acknowledgements

We thank K. Hantke for valuable comments on the manuscript.

