## Supplementary Material for "Curcumin-mediated photodynamic inactivation of *Escherichia coli, Pseudomonas fluorescens*, and *Candida auris*"

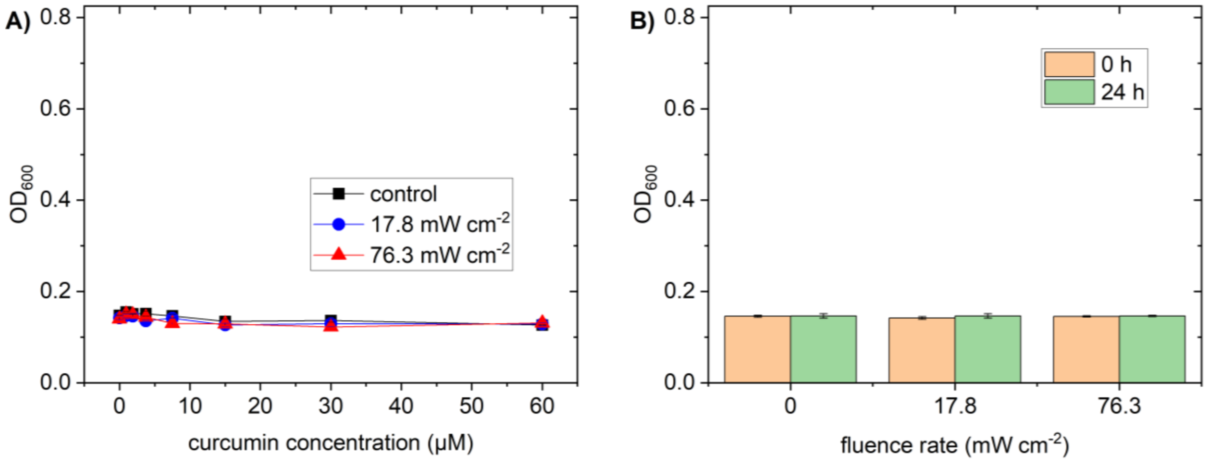


**Figure S1**. Optical density ($OD_{600}$) of the E. *coli* samples and controls shown in Figure 1A. A) $OD_{600}$ of samples and dark control before incubation (0 h). B) $OD_{600}$ of negative controls without bacteria. Error bars indicate the standard deviation (n = 3).

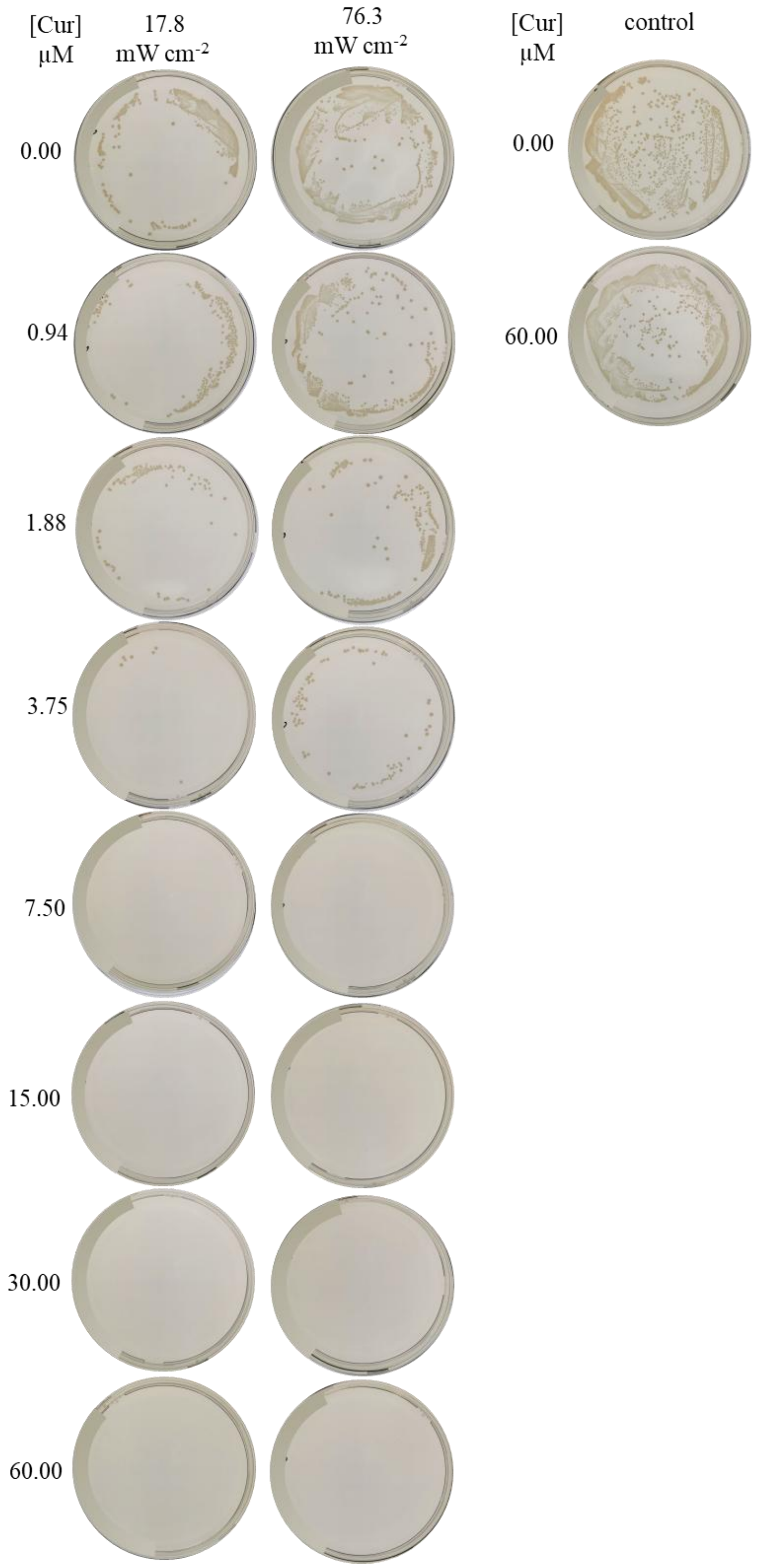


**Figure S2**. Agar plates of *E. coli* colonies after curcumin-mediated aPDT using 420 nm light with a fluence rate of 17.8 mW $cm^{-2}$ and 76.3 mW $cm^{-2}$, respectively, at different curcumin concentrations. The agar plates were incubated for 24 h at 37 $^{o}C$. Controls were incubated in the dark.

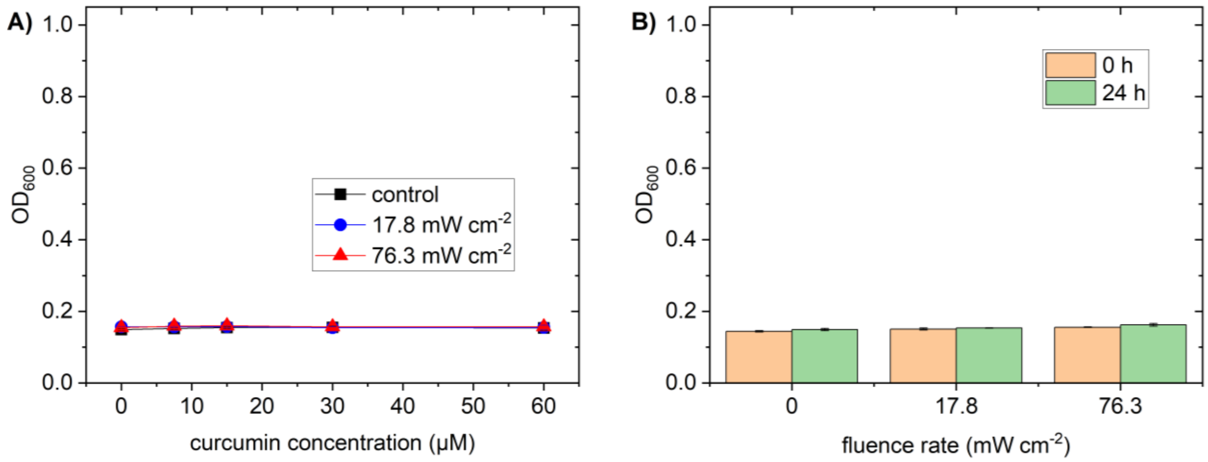


**Figure S3**. Optical density (OD$_{600}$) of the *P. fluorescens* samples and controls shown in Figure 2A. A) OD$_{600}$ of samples and dark control before incubation (0 h). B) OD$_{600}$ of negative controls without bacteria. Error bars indicate the standard deviation (n = 3).

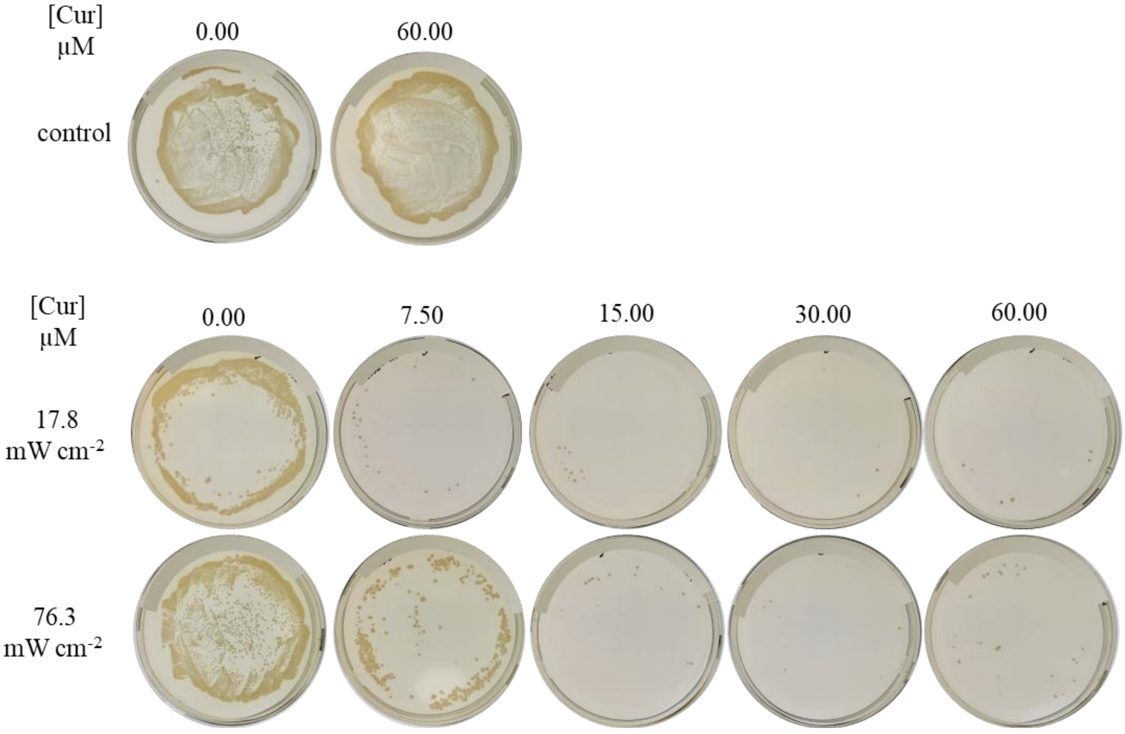


**Figure S4**. Agar plates of *P. fluorescens* colonies after curcumin-mediated aPDT using 420 nm light with a fluence rate of 17.8 mW cm$^{-2}$ and 76.3 mW cm$^{-2}$, respectively, at different curcumin concentrations. The agar plates were incubated for 48 h at 30 $^{o}$C. Controls were incubated in the dark.

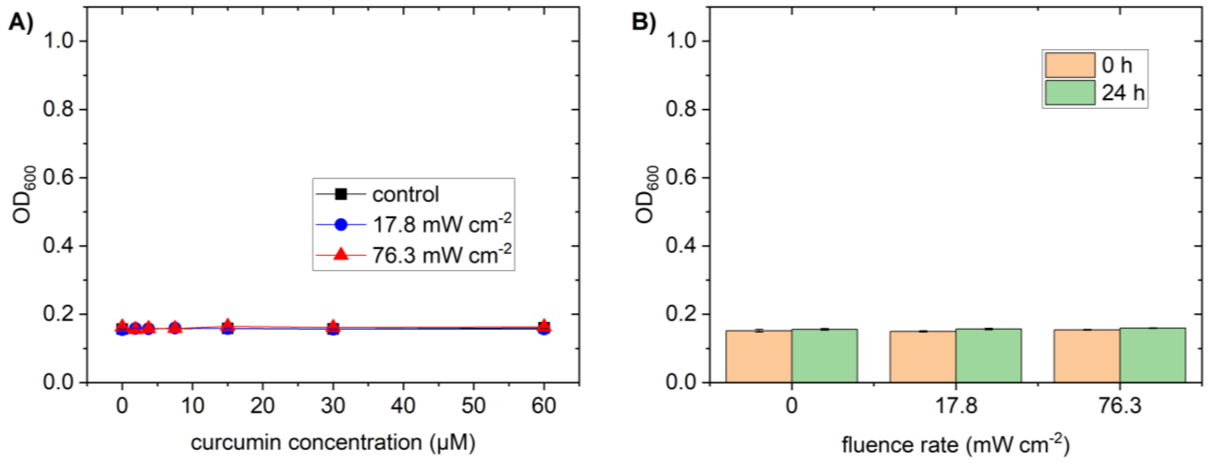


**Figure S5**. Optical density ($OD_{600}$) of the *C. auris* samples and controls shown in Figure 3A. A) $OD_{600}$ of samples and dark control before incubation (0 h). B) $OD_{600}$ of negative controls without fungi. Error bars indicate the standard deviation (n = 3).

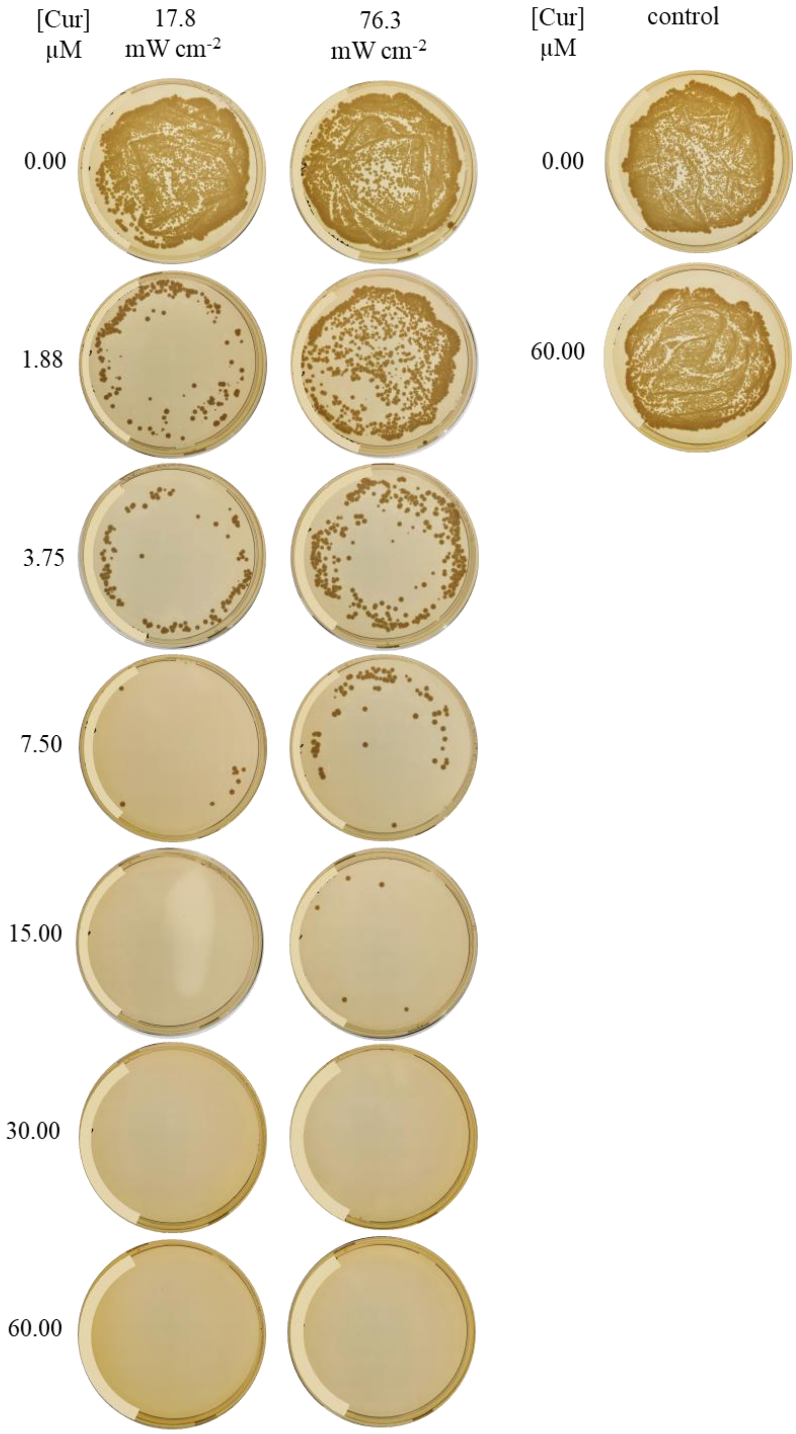


**Figure S6**. Agar plates of *C. auris* colonies after curcumin-mediated aPDT using 420 nm light with a fluence rate of 17.8 mW cm$^{-2}$ and 76.3 mW cm$^{-2}$, respectively, at different curcumin concentrations. The agar plates were incubated for 24 h at 30 °C. Controls were incubated in the dark.
